# Pho-Tip: one-pot dephosphorylation for rapid and sensitive analysis of DIA phosphoproteomics data

**DOI:** 10.1101/2025.11.17.687597

**Authors:** Katharina D. Faisst, Kate Lau, Ludwig R. Sinn, Lukasz Szyrwiel, Vadim Demichev

**Affiliations:** Quantitative Proteomics laboratory, Charité – Universitätsmedizin Berlin, Germany

## Abstract

Recent advances in instrumentation and data processing have transformed data-independent acquisition (DIA) proteomics into a reliable technology for quantitative profiling of post-translational modifications. However, analysis of DIA phosphoproteomics data is challenging due to the large search space, wherein all combinations of phosphosites on a peptide need to be considered. Current approaches therefore face significant hurdles in detecting low-abundant phosphorylated peptides, in particular when working with low sample amounts. Here we introduce Pho-Tip, a lossless one-pot dephosphorylation strategy. We show that Pho-Tip enables comprehensive mapping of phosphorylated peptide sequences, facilitating streamlined creation of experiment-focused *in silico* predicted spectral libraries and thus rapid and sensitive analysis of DIA phosphoproteomics experiments.

## Introduction

Data-independent acquisition (DIA) proteomics has gained significant popularity in recent years, promoted by step change improvements in mass spectrometry instrumentation and data analysis software algorithms^1,2^. DIA has also been established for the analysis of post-translational modifications (PTMs), including phosphorylation^3^. At the same time, the accepted gold standard approach to phosphoproteomics involves spectral library creation via deep offline fractionation-based analysis of a pooled sample. Given that this process is laborious and is not guaranteed to enable full coverage of detectable phosphosites throughout the experiment, library-free analysis of DIA phosphoproteomics data has emerged as a viable alternative, enabled by the advances in DIA data processing software^4,5^.

This approach, however, results in a large search space, as a single peptide sequence can give rise to multiple possible combinations of occupied phosphosites. The large search space reduces sensitivity, since the DIA software necessarily has to impose stricter quality requirements on peptide spectrum matches in order to report them as confidently identified. Further, the analysis time grows along with the search space, presenting a considerable computational burden, this being particularly relevant in view of the increasing proportion of large-scale proteomics experiments among the applications of DIA^6,7^.

To address the search space problem of DIA phosphoproteomics, we introduce Pho-Tip, a one-pot dephosphorylation strategy that leverages phosphopeptide enrichment followed by alkaline phosphatase treatment on EvoTips (Evosep) and liquid chromatography coupled to mass spectrometry (LC-MS) analysis. Subsequent search against the full sequence database provides comprehensive identification of any amino acid sequences which may bear a phosphorylation. These sequences can then serve as the basis for predicting compact phosphopeptide-containing spectral libraries for the sensitive analysis of phosphoproteomics experiments. Here, we demonstrate that this strategy outperforms the conventional search against the full sequence database in terms of speed and coverage.

## Results

In the past, dephosphorylation has already been considered for the purpose of enhancing phosphopeptide detection, in combination with data-dependent acquisition (DDA) mass spectrometry. However, the inherent stochastic nature of DDA tends to result in a high missing value rate, and, consequently, early experiments were not able to achieve comprehensive phosphoproteome coverage^8,9^. We speculated that high-sensitivity DIA proteomics, in contrast, can comprehensively capture the phosphoproteome via the analysis of a dephosphorylated sample.

To confirm this, we have enriched phosphopeptides from a HeLa cell line tryptic proteome digest using TiO_2_ columns (Thermo Fisher Scientific), followed by treatment with calf intestinal alkaline phosphatase (CIP) or a mock treatment. The samples were then analysed on a ZenoTOF 7600 mass spectrometer (SCIEX), using a 20-min active chromatographic gradient (Methods). First, the data were searched with DIA-NN 2.0.2 against the human sequence database, with up to three phosphorylations enabled, confirming that the CIP treatment successfully removes the vast majority of peptide phosphorylations (Supp. Fig. 1). We then searched the CIP-treated sample against a database devoid of any variable peptide modification, resulting in a compact search space size for maximum identification sensitivity. In support of our hypothesis, we observed that the CIP-treated sample allowed the detection of 94% of amino acid sequences identified as phosphorylated in the mock-treated sample (n = 7,119). Moreover, the total number of detected peptide sequences, regardless of the phosphorylation status, increased from 15,014 to 27,962 upon CIP treatment, hinting at potentially higher sensitivity of peptide detection in their dephosphorylated form. To investigate the possible reasons for the increased sensitivity beyond the search space reduction, we compared the MS1 signal attributed to each amino acid sequence in its phosphorylated form in the mock-treated sample to the respective MS1 signal detected in the CIP-treated sample. Jointly detected amino acid sequences devoid of any phosphosites served as loading control. This analysis revealed a minor increase in detected signal following dephosphorylation (Supp. Fig. 2).

Next, we established Pho-Tip, a streamlined pipeline that enables to easily obtain the set of phosphopeptide amino acid sequences for the vast majority of phosphoproteomics applications. For this, we coupled enrichment using MagReSyn® Zr-IMAC HP magnetic beads (ReSyn Biosciences), previously established as compatible with a wide range of peptide loads^10^, to peptide dephosphorylation on EvoTips. Recent work demonstrated how EvoTips can be used for lossless sample preparation via on-tip reactions^11^, causing us to speculate that we can perform dephosphorylation on-tip too. To evaluate the possible losses associated with it, we have performed phosphopeptide enrichment based on 500µg of yeast (*S. cerevisiae*) tryptic proteome digest and loaded the enriched phosphopeptides on EvoTips, subsequently either spinning them down immediately (*i*.*e*., “mock treatment”) or first treating with CIP for 2 h (Methods), prior to analysis on Evosep One (Evosep) coupled to a timsTOF Ultra (Bruker) mass spectrometer using a 60 SPD (samples per day) method (Methods). There was no significant difference in the detected intensities of peptides that do not contain serines, threonines or tyrosines, confirming that the CIP treatment on-tip can be considered lossless (Supp. Fig. 3). Analysing the sequence coverage obtained with the CIP-treated sample, we again observed 94% of sequences detected as phosphorylated in the mock-treated sample.

We then proceeded to investigate how searching phosphopeptide data (that is, generated from samples without CIP treatment) against sequences exclusively identified in CIP-treated samples affects the numbers of detected phosphopeptides, compared to searching against the fully predicted sequence database of the respective species (Fig. 1a; human or yeast). Here, setting the maximum number of variable modifications to 3, allowing 2 missed cleavages and precursor charges 2-3 resulted in spectral libraries of the size ∼896k precursors (human) and ∼791k precursors (yeast), compared to ∼24.3M and ∼10.8M precursors, respectively, in case of the full sequence search space. Searching the phosphopeptide samples, we observed that CIP-based predicted libraries resulted in identification of about +15% on average for the human sample and about +2% for the Pho-Tip-processed yeast sample. The gains appeared distributed across the whole range of precursor masses and retention times (Supp. Fig. 4).

**Fig. 1.**
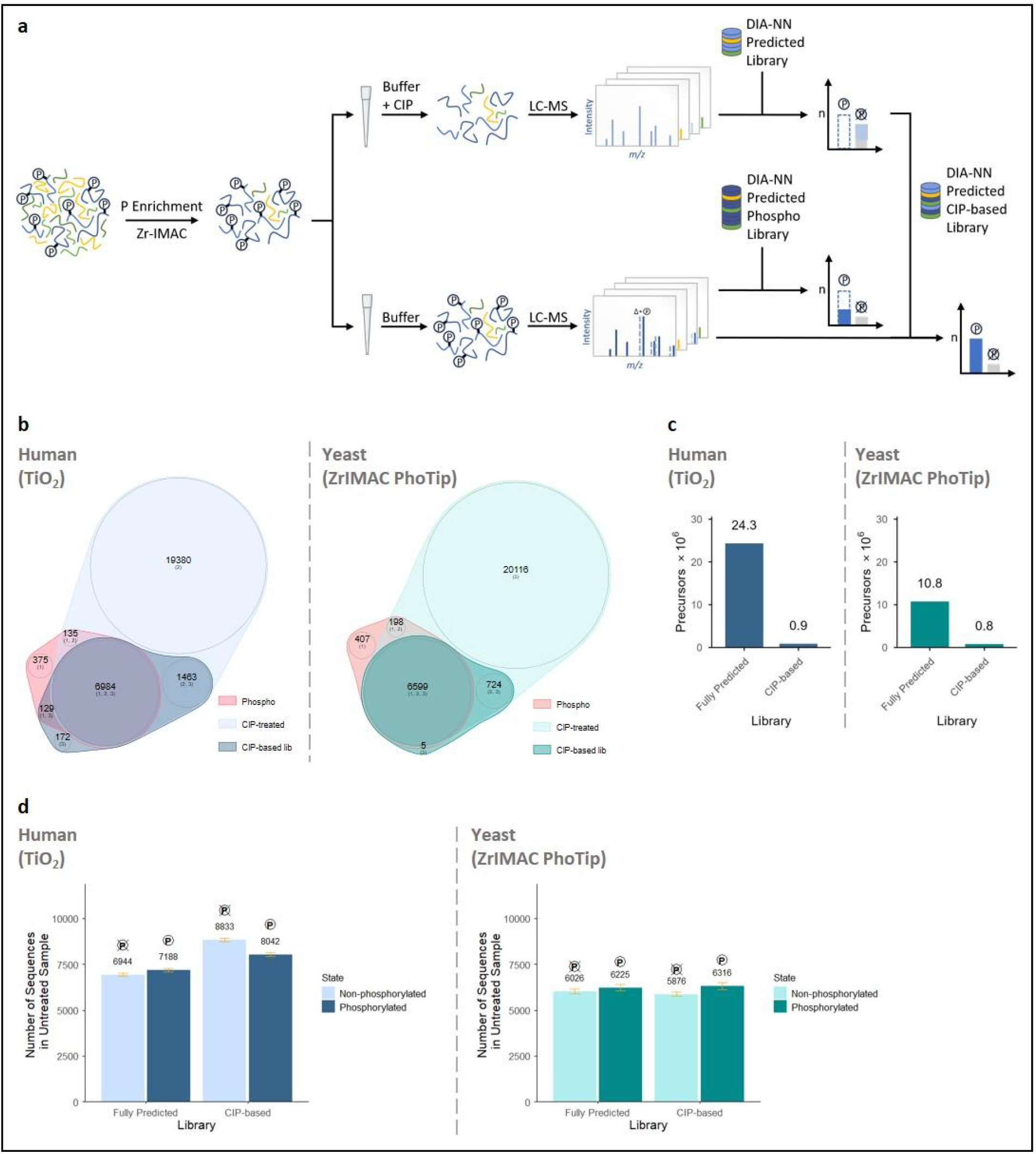
Pho-Tip improves phosphoproteomics sensitivity and boosts data analysis speed. **a)** Pho-Tip experimental workflow including phosphopeptide enrichment and CIP-treatment on-tip, analysis on Evosep One coupled to timsTOF Ultra and searching of phosphopeptide samples against exclusively sequences found in CIP-treated samples, compared to searching against the full sequence database. **b**) Overlap of identified phosphorylated peptide sequences (Phospho) with CIP-treated (CIP-treated) and mock-treated sample analysed using the CIP-based library (CIP-based library), separately for TiO_2_ as well as Zr-IMAC coupled to Pho-Tip. **c**) Precursor counts for the full sequence database-based and CIP-based in silico predicted libraries, reflecting the analysis speed. **d**) Number of identified sequences split by phosphorylation state (phosphorylated vs non-phosphorylated) comparing full sequence database-based and CIP-based library search, separately for TiO_2_ as well as Zr-IMAC coupled to Pho-Tip.

We further investigated if Pho-Tip can be used directly in quantitative proteomics experiments. While in many cases knowing the exact phosphosite location is essential in the context of the experiment, the numbers of sites confidently localised are typically low, often fold-change lower than the numbers of detected phosphopeptides. This leads to the 75% confidence threshold, as reported by the analysis software, often being used as the basis of phosphosite identification and subsequent quantification, but even with such a loose filtering many sites are being discarded. We note that it is common to perform preliminary discovery screens that are then followed by validation based on selected samples of interest, at a lower scale and possibly higher sample amounts and longer chromatographic gradients, possibly combined with sensitive parallel reaction monitoring (PRM), resulting in improved data quality. We speculate that in some cases discovery screens may not require resolution down to a specific site, but rather could focus on detection of differentially regulated peptide species, with subsequent investigation uncovering the exact sites involved. Such a design would benefit from maximum possible sensitivity, making Pho-Tip an ideal building block for a candidate screening method. Here, we note that coefficients of variation (CVs) observed for peptides in Pho-Tip or TiO_2_-enriched CIP-treated samples data are comparable to those observed without CIP treatment (Supp. Fig. 5).

## Discussion

In this work, we tackle the computational challenge of phosphoproteomics: a large search space when considering all possible phosphopeptides that results in decreased sensitivity for phosphopeptide identification as well as a demand for large amounts of computational resources. While the creation of spectral libraries via offline fractionation of a pooled sample is an option, it may be too labour intensive to perform it for each project, and it further requires significant sample amounts to be effective. In addition, peptides with rare phosphosite occupancy configurations may be missed using the pooled library approach.

We propose a different, ‘biochemical’, solution: the search space is reduced in-sample, by enzymatically dephosphorylating all phosphopeptides, wherein multiple peptide species are converted into a single unmodified peptide sequence. With Pho-Tip being a lossless reaction, as well as given the improved detectability of unmodified peptides compared to phosphopeptides, sensitivity for phosphopeptide sequence detection is maximised. The identified peptide sequences then allow the creation of experiment-focused *in silico*-predicted spectral libraries that achieve fast and comprehensive analysis of regular, “CIP-untreated”, phosphopeptide-enriched samples.

We note that the benefits of Pho-Tip may extend beyond just speeding up the analysis and increasing its sensitivity. In fact, in Pho-Tip each phosphopeptide sequence is detected twice, in modified and unmodified form. It is natural to assume that such repeated detection would result in a lower false discovery rate at the same q-value reported by the software, possibly allowing to filter the data at a q-value threshold less stringent than the commonly accepted 1%. We will leave it for future investigations to assess the magnitude of this effect with a suitable experiment design.

We envision that Pho-Tip can be used in a number of distinct ways as part of a phosphoproteomics workflow with the option for automation. For example, a pooled sample or a subset of samples may be analysed via Pho-Tip, to create the spectral library. Alternatively, we speculate that Pho-Tip can prove to be an excellent quantitative discovery tool in fast phosphoproteomics screens, used separately or in conjunction with regular phosphoproteomics sample acquisition. In this context, differentially expressed peptide sequences identified with Pho-Tip may subsequently be validated to pinpoint the involved phosphosites.

In conclusion, we expect Pho-Tip to make DIA phosphoproteomics more accessible, abolishing the need for the conventional spectra library creation.

## Methods

### Sample Preparation - Yeast

Yeast strain BY4741ki^12^ was cultured in Yeast Extract Peptone Dextrose (YPD) medium at 30°C, 270 rpm. The cultures were harvested by centrifugation with 3,000 x g for 15 min at 4°C. Pellets were washed with 1x phosphate buffered saline (PBS) containing 0.8 mM phenylmethylsulfonyl fluoride (PMSF), aliquoted and stored at -80°C. Cells were thawed, washed with ice-cold MilliQ water and pretreated with 0.2 M sodium hydroxide (NaOH) for 10 min on ice. After pelleting, cells were resuspended in lysis buffer (7M Urea, 0.1M ammonium bicarbonate (ABC)) containing PhosphataseArrest™ I (G-Biosciences, St. Louis, MO, USA), mixed with 0.5 mm glass beads, and subjected to lysis by bead-beating using the SPEX CertiPrep™ Geno/Grinder 2 (SPEX SamplePrep, Metuchen, NJ, USA) at 1,500 rpm for four 5 min cycles, with cooling on ice between cycles. The samples were centrifuged at 20,000 x g for 15 min at 4°C, and the supernatants were collected, and centrifuged again at 20,000 x g for 1h at 4°C. The final supernatant was used for protein concentration determination using the Pierce™ BCA Protein Assay Kit (Thermo Fisher Scientific, Waltham, MA, USA).

For disulfide reduction, the sample was incubated with 5 mM dithiothreitol (DTT) at RT for 30 min, then kept on ice for 5 min. Alkylation was performed by adding 10 mM iodoacetamide (IAA), followed by incubation in the dark for 20 min. The reaction was quenched with an additional 5 mM DTT, and incubated in the dark for 10 min. The cell lysate was diluted 1:5 with 0.1 M ABC, and trypsin was added at an estimated 1:100 (m/m) enzyme-to-protein ratio. The mixture was vortexed, centrifuged, and incubated overnight at 37°C. Digestion was stopped by acidifying with trifluoroacetic acid TFA to pH 2-3. After 15 min incubation at RT, the samples were centrifuged at 4°C, and the supernatants were collected.

### Sample Preparation - HeLa

HeLa cells were cultured in Dulbecco’s Modified Eagle’s Medium (DMEM) medium (500 mL) containing L-glutamine, supplemented with 10% fetal calf serum (FCS) and 1% Penicillin-Streptomycin. Cells were maintained at 37°C in a humidified atmosphere with 5% CO_2_. The cultures were washed with PBS and harvested using ethylenediaminetetraacetic acid (EDTA). Cells were then washed with PBS containing 0.8 mM PMSF, aliquoted, pelleted (5x10^6 cells per pellet) and stored at -80°C. Cells were thawed, washed with ice-cold MilliQ water and afterwards resuspended in lysis buffer (0.1 M TRIS–HCl, 8 M Urea) containing PhosphataseArrest™ I. The sample was vortexed and sonicated in an ultrasonic bath for 30 s. Homogenization was performed using a 26G needle. Subsequently, benzonase was added, and the mixture was incubated at RT for 15 min. After incubation, the mixture was centrifuged at 14,800 rpm for 60 min, and the supernatant was collected. Protein concentration was then determined using the Pierce™ 660 nm Protein Assay (Thermo Fisher Scientific, Waltham, MA, USA).

For disulfide reduction, the sample was incubated with 10 mM DTT at RT for 30 min, then kept on ice for 5 min. Alkylation was performed by adding 20 mM IAA, followed by incubation in the dark for 20 min. The reaction was quenched with an additional 10 mM DTT, and incubated in the dark for 10 min. The cell lysate was diluted 1:5 with 0.1 M ABC, and trypsin was added at a 1:100 enzyme-to-protein ratio. The mixture was vortexed, centrifuged, and incubated overnight at 37°C. Digestion was stopped by acidifying with TFA to pH 2-3. After 15 min incubation at RT, the samples were centrifuged at 4°C, and the supernatants were collected.

The yeast and HeLa samples were desalted using STAGE-Tips according to the protocol.^13^ STAGE-Tips were activated with methanol (MeOH), and washed with 60% (v/v) acetonitrile (ACN) and 0.1% (v/v) TFA. Samples were loaded onto the tips, and washed with 0.1% TFA. Peptides were eluted with 60% ACN. The eluates were dried using the Eppendorf^®^ Concentrator Plus (Eppendorf SE, Hamburg, Germany) at 45°C, and reconstituted in 2% ACN, 0.1% TFA. Peptide concentrations were determined using the Implen NanoPhotometer® N60/N50 (Implen GmbH, München, Germany). Samples were stored at -80°C.

### Phosphopeptide Enrichment using the High-Select™ TiO_2_ Phosphopeptide Enrichment Kit and Phosphatase Reaction - HeLa

Phosphopeptide enrichment was performed with the High-Select™ TiO_2_ Phosphopeptide Enrichment Kit (Thermo Fisher Scientific, Waltham, MA, USA) according to the manufacturer’s instructions. In brief, dried peptide pellets (3 replicates per condition) were resuspended in binding/equilibration buffer. Columns were first washed before equilibration, both with centrifugation at 3000×g for 2 min. The sample was applied and loaded at 1000×g for 5 min. The flow-through was reapplied to ensure full binding, with the same centrifugation and retention of the flowthrough. Afterwards, the column was washed by adding first Binding/Equilibration and then wash buffer, followed by centrifugation at 3000×g for 2 min. These two washing steps were repeated, and the column was then washed with LC-MS grade water. To elute the phosphopeptides, 25 µl elution buffer was added. The column was centrifuged at 1000×g for 5 min, and the elution step was repeated with 25 µl elution buffer. The eluates were dried using the Eppendorf^®^ Concentrator Plus at 45°C, and reconstituted in 2% ACN, 0.1% TFA. Peptide concentrations were determined using Nanodrop.

For phosphatase reaction the peptide lysate was first mixed with a pre-prepared phosphatase reaction buffer, then divided equally between two tubes. For CIP-treated samples, CIP was added to one of the tubes while the other tube was considered mock-treated. The reaction was carried out by incubating both mock- and CIP-treated samples at 37°C for 2 h. Following incubation, samples were quenched by adding TFA to a final concentration of 0.5%.

The samples were again desalted using STAGE-Tips according to the protocol. STAGE-Tips were activated with MeOH, and washed with 60% ACN and 0.1% TFA. Samples were loaded onto the tips, and washed afterwards with 0.1% TFA. Peptides were eluted with 60% ACN. The eluates were dried using the Eppendorf^®^ Concentrator Plus at 45°C, and reconstituted in 2% ACN, 0.1% TFA. Peptide concentrations were determined using Nanodrop. Samples were stored at -80°C.

### Phosphopeptide Enrichment with MagReSyn® Zr-IMAC HP Beads and Phosphatase Reaction - Yeast

Phosphopeptide enrichment of yeast proteome digests was performed using MagReSyn® Zr-IMAC HP beads according to the manufacturer’s instructions. Dried peptide pellets were resuspended in binding buffer. MagReSyn^®^ Zr-IMAC HP beads were equilibrated by resuspending in 500 µl binding buffer for three cycles. The equilibrated beads (5 µl) were distributed into wells and 30 µg of resuspended peptides were added (6 replicates per condition). The mixture was incubated mixing on a Eppendorf ThermoMixer^®^ C (Eppendorf SE, Hamburg, Germany) at 750 rpm for 20 min. Beads were washed sequentially with binding buffer, Wash Solvent 1 (80% ACN + 1% TFA), and Wash Solvent 2 (10% ACN + 0.2% TFA). Phosphopeptides were eluted from the beads using 50 µl of 0.5%, 1%, and 2% NH_4_OH solutions, pooled, and the pH was adjusted to 9-10 with 10% TFA. For the following dephosphorylation reaction, 10x CIP buffer was added to achieve a final concentration of 1x.

EvoTips were prepared by activation and equilibration according to the manufacturer’s protocol. Peptide samples (30 ng) were loaded per EvoTip. For mock-treated samples, the loaded EvoTips were spun down immediately without further treatment. For CIP-treated samples, CIP was added to the loaded EvoTips, followed by incubation at 37°C for 2 h. After incubation, CIP-treated samples were quenched with 20% formic acid (FA). All EvoTips were subsequently washed with 0.1% FA and then loaded with 100 µl of 0.1% FA in preparation for analysis.

### LC-MS/MS Analysis - TiO_2_ HeLa

Samples were acquired in data-independent acquisition (DIA)/ Zeno-SWATH mode, on a ZenoTOF 7600 mass spectrometer (SCIEX, Toronto, Canada) coupled online to an Acquity M-Class UPLC system (Waters, Milford, MA, USA). In each measurement we injected 200 ng of sample onto a reversed-phase chromatography nanoEase M/Z HSS T3 column (100 Å, 1.8 µm, 0.3 x 150 mm) with buffers A (0.1% FA) and B (acetonitrile with 0.1% FA) ramping from 3 to 40% buffer B at a flow rate of 5 µl/min and at 35°C using a 30 min total gradient as described in Wang *et al*. 2022^14^.

The mass spectrometric method was similar to the one employed by Wang and colleagues^14^ with the following exceptions: 85 variable isolation windows between 4 and 8 Th were defined. The spray voltage was set to 5000 V and a scheduled ionization between minutes 2.5 and 22 was used.

### LC-MS/MS Analysis - ZrIMAC Yeast

LC-MS was performed using an Evosep One system coupled to a TimsTOF ULTRA mass spectrometer equipped with a Captive Spray II ion source. Peptide separation was conducted on an 8 cm × 150 µm ID Performance Column with 1.5 µm particle size, maintained at 40°C. A standard 60 SPD Evosep method was applied, utilizing a 21-min gradient for a total sample-to-sample time of 24 min. Solvent A consisted of 0.1% FA, and Solvent B was ACN with 0.1% FA (Optima LCMS grade, Thermo Fisher Scientific, Waltham, MA, USA).

Data were acquired using a dia-PASEF scheme, optimized for a cycle time of 1.18 s. The isolation window scheme spanned m/z 388−1168 and 1/K0 0.67−1.28, covering most charge 2 precursor ions, using 28 × 27.8 Th windows. The acquisition utilized 100 ms accumulation and ramp times. The mass spectrometer was operated in “high sensitivity detection” mode, optimized for low sample amounts.

The predefined dia-PASEF window scheme was selected to balance sensitivity and coverage, though some bias in precursor ion sampling may remain inherent to the acquisition set-up.

### Raw Data Analysis - TiO_2_ HeLa and ZrIMAC Yeast

DIA-NN 2.0.2 was used to analyse the experiment, the full settings are reflected by the logs deposited to the PRIDE repository. Briefly, mass accuracy was set to 15 ppm (timsTOF) or 20 ppm (ZenoTOF); MS1 accuracy was 15ppm (timsTOF) or 12 ppm (ZenoTOF). The precursor charge range was set to 2 to 3, the precursor’s m/z range was set to 400 to 1200.

### Spectral Library Generation

To generate a CIP-based library, CIP-treated samples were analysed without specifying phosphorylation as a variable modification, with Scoring set to Generic and FDR cutoff set to 5%. The generated .skyline.speclib file was then used to create a predicted phospho library, with it being added instead of a FASTA database using the “Add FASTA” button in DIA-NN, and with --cut provided in Additional options field to disable enzymatic digest of the library peptide sequences. This resulted in a library that contains all possible phosphorylation states of peptide sequences that were detected in CIP-treated samples. For all other analyses, Scoring was set to Proteoforms. When generating predicted libraries, the maximum number of missed cleavages was set to 2, precursor charge range was 2 to 3 and precursor m/z range was 400 to 1200. Where applicable, phosphorylation was enabled as variable modification and the maximum number of variable modifications was set to 3. This method generates a tailored spectral library, focusing on phosphorylated peptides from the dataset, instead of relying on a generic protein database.

## Data Availability

The MS proteomics data have been deposited to the ProteomeXchange Consortium via the PRIDE^15^ partner repository with the dataset identifier PXD070420.

## Acknowledgements

This work is funded by the German Ministry of Education and Research (BMBF), as part of the National Research Node “Mass spectrometry in Systems Medicine” (MSCoreSys), under grant agreement 161L0221.

## Supplement

**Supp. Fig. 1.**
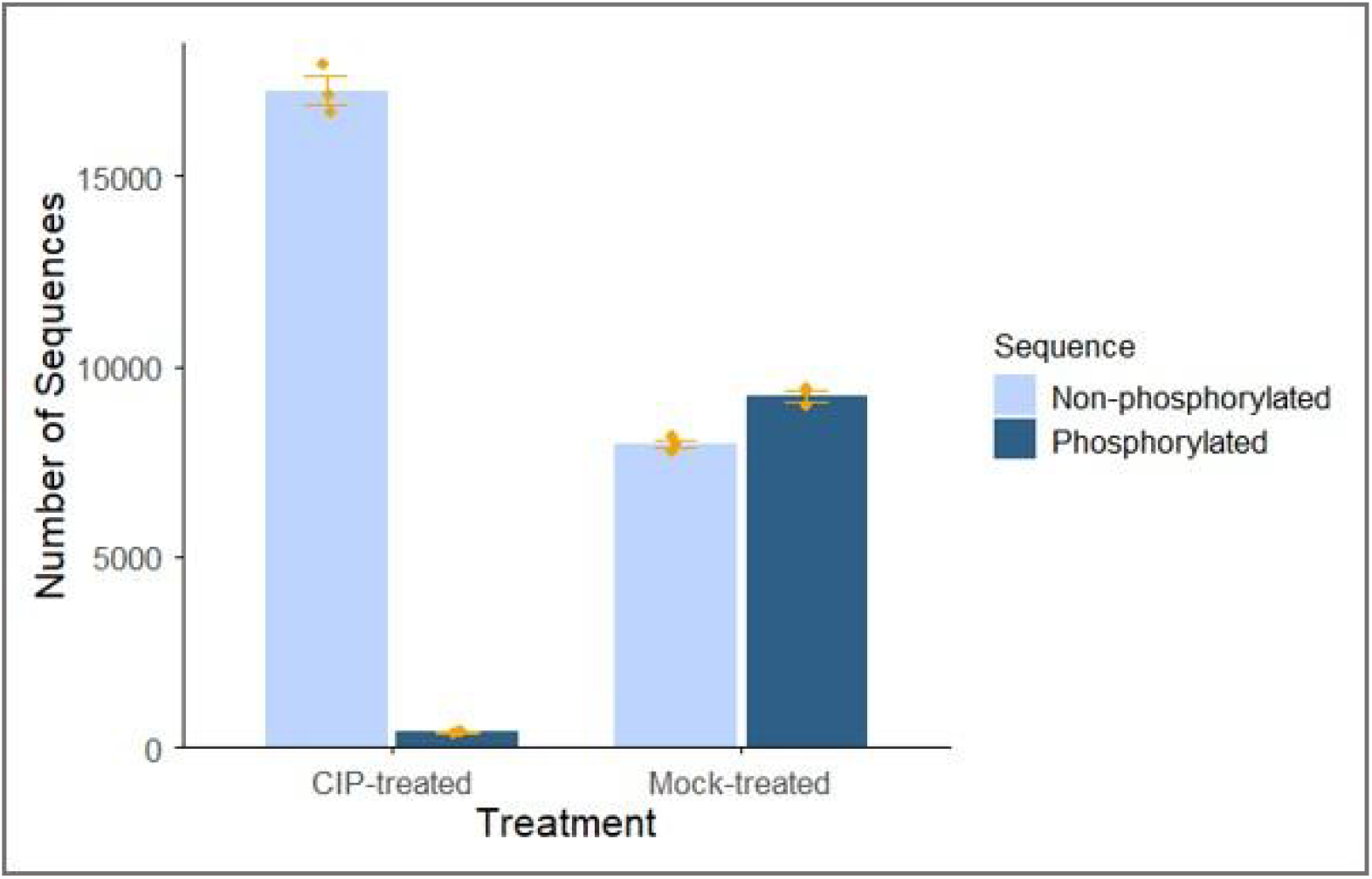
CIP-treatment efficiently desphosphorylates peptides. Number of phosphorylated and non-phosphorylated precursors in mock- and CIP-treated samples in human TiO_2_-enriched samples. Group means are shown, error bars indicate standard deviation.

**Supp. Fig. 2.**
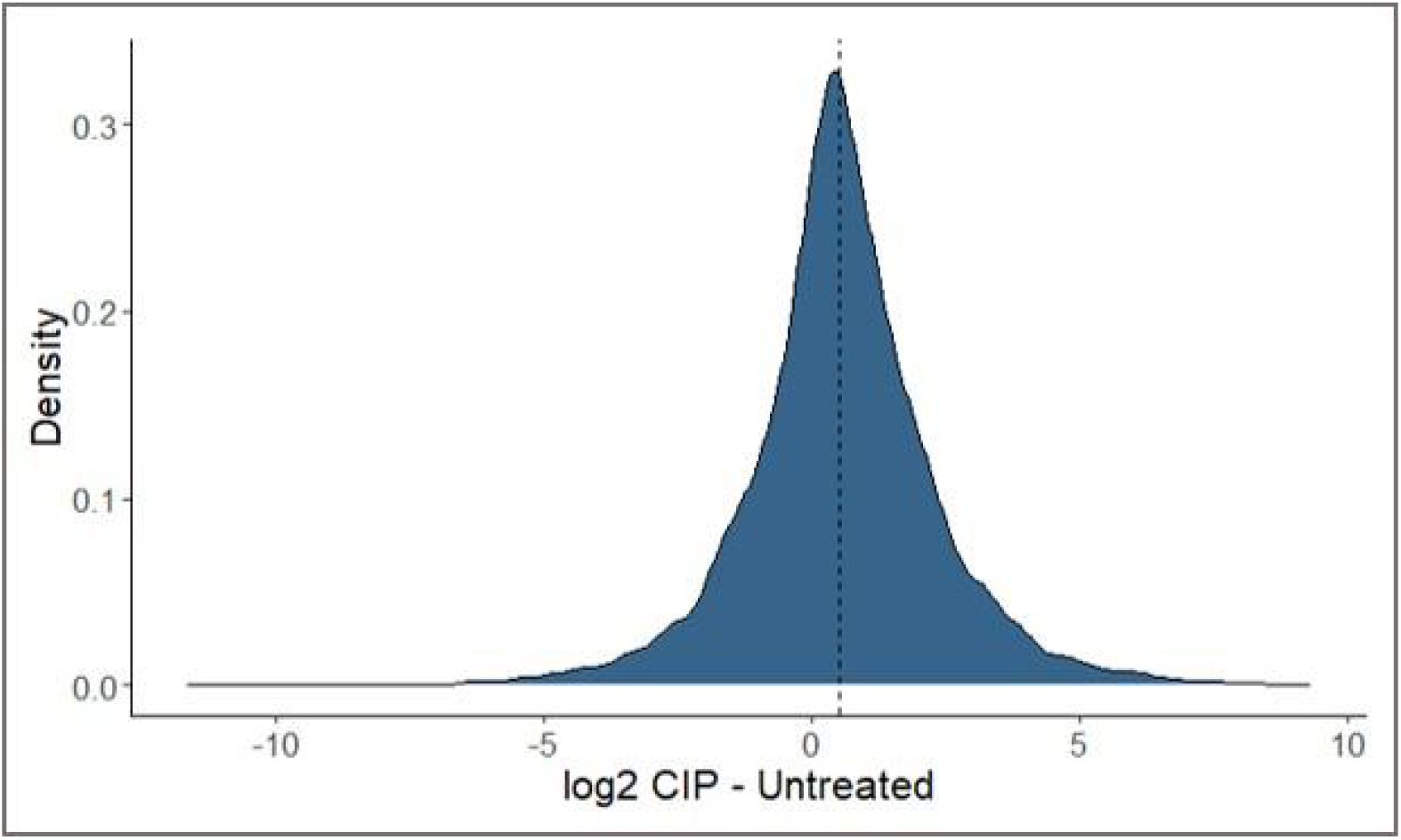
Dephosphorylation improves the detectability of peptides. Density distributions of log2-transformed integrated MS1 signal differences between unphosphorylated peptide sequences detected in CIP-treated samples and their phosphorylated counterparts in mock-treated samples for human TiO_2_-enriched samples. Intensities of the latter were aggregated, for each stripped sequence, using the maximum value across matching precursors. Data were normalized to non-phosphorylatable peptides (those lacking Ser/Thr/Tyr residues) in the respective samples. The median value (0.31) is indicated with a dotted line.

**Supp. Fig. 3.**
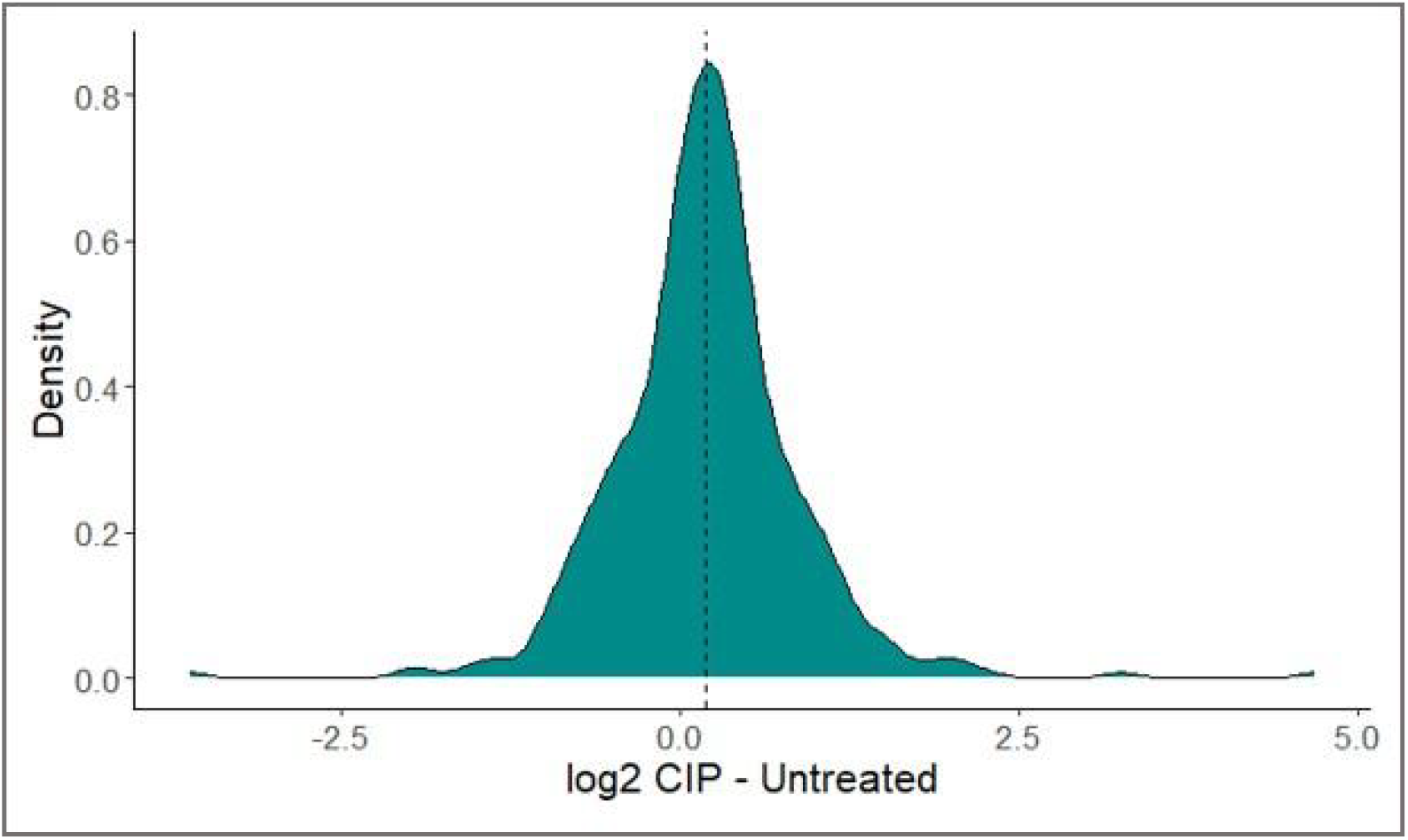
Pho-Tip results in lossless dephosphorylation. Density distributions of log2-transformed integrated MS1 signal differences between unphosphorylated peptide sequences detected in CIP-treated Pho-Tip samples and their phosphorylated counterparts in untreated samples. Non-phosphorylatable peptides (those lacking Ser/Thr/Tyr residues) are shown. Intensities were aggregated using the maximum values. The median value (0.19) is indicated with a dotted line.

**Supp. Fig. 4.**
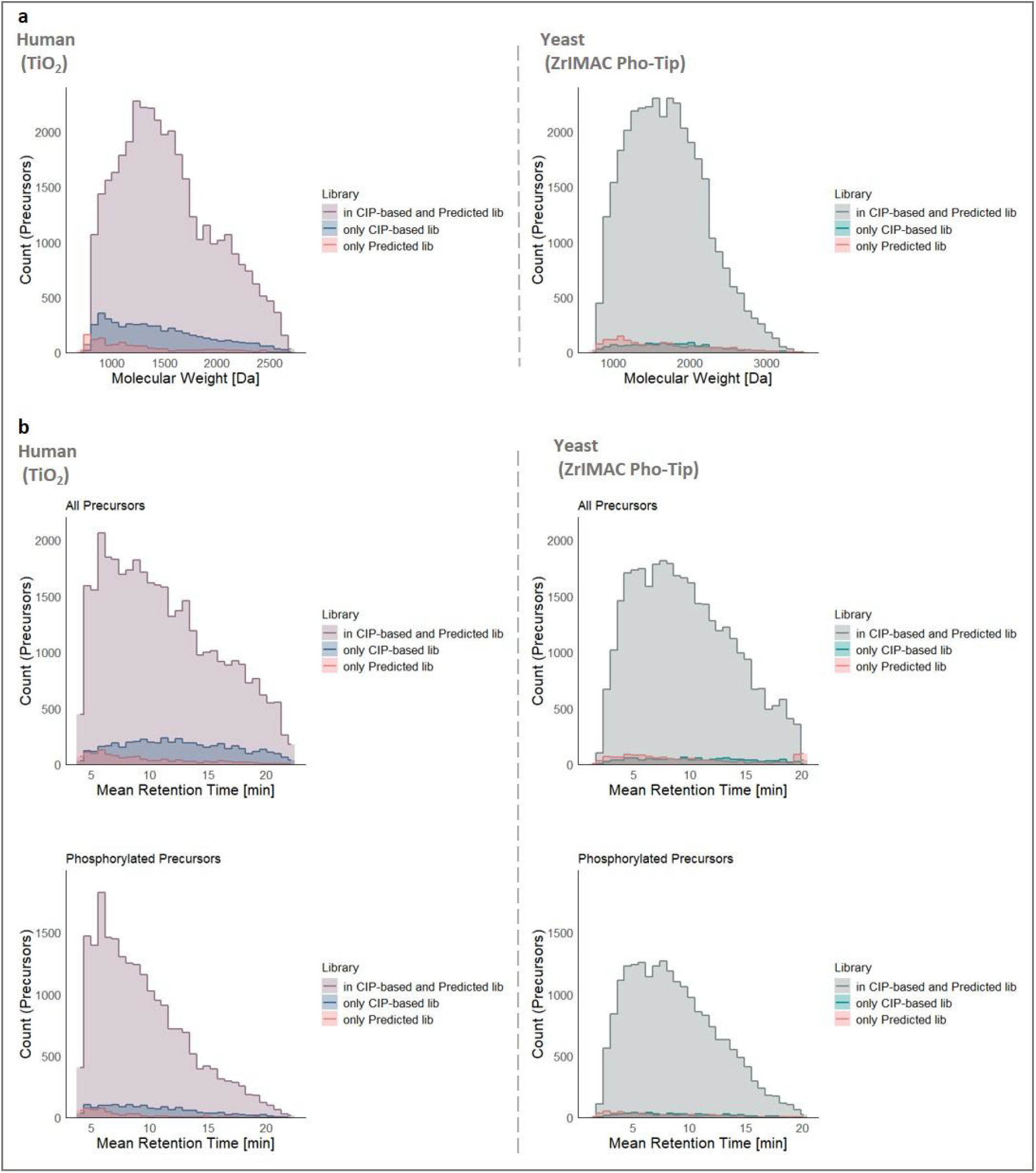
Precursor properties distributions depending on the search mode. **a)** Distribution of molecular weight of precursors for human TiO_2_-enriched and yeast Zr-IMAC Pho-Tip samples when searching with each library. **b)** Same for retention times.

**Supp. Fig. 5.**
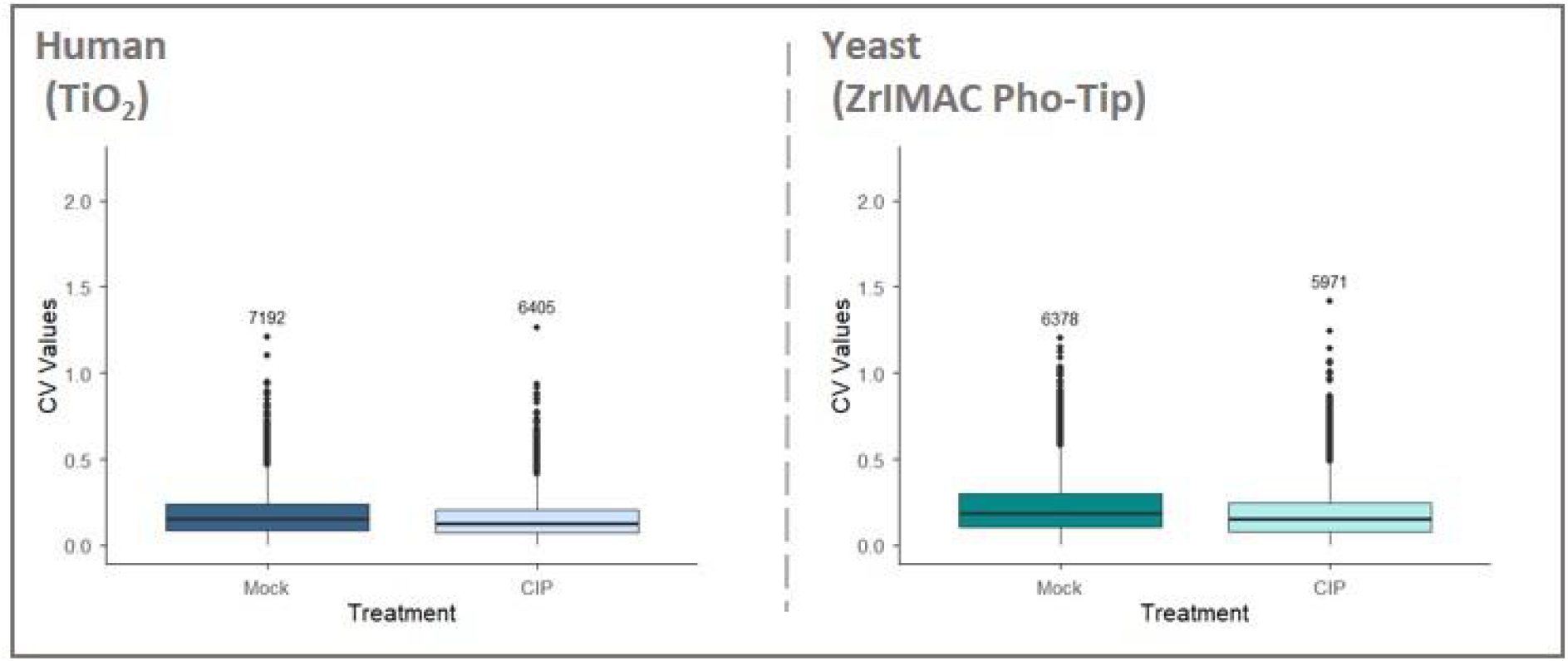
The effect of dephosphorylation on quantitative precision. Precursor-level CV values for jointly detected precursors by treatment, separately for human TiO_2_-enriched and yeast Zr-IMAC Pho-Tip samples. The boxes show the interquartile range (IQR) and the whiskers expand 1.5x the IQR.

